# Variant effect predictors are shaping the ClinVar classifications used to train and evaluate them

**DOI:** 10.64898/2026.10.24.761512

**Authors:** Minyu Tao, Joseph A. Marsh

## Abstract

**Summary:** Variant effect predictors (VEPs) are playing an increasingly influential role in clinical variant classification. Following recent calibration efforts and the development of points-based classification standards, computational predictions can now carry enough weight to move variants out of uncertain significance with little additional evidence. While clinically valuable, this increases the potential for classification-level circularity, where VEP predictions help determine the clinical classifications later used to train, benchmark and calibrate VEPs. Here, we show that computational evidence is already leaving a detectable signature in ClinVar. Among rare missense variants from a major submitter, likely benign classifications shifted markedly towards more benign scores and show a sharp step change at the calibrated moderate-evidence threshold of BayesDel_noAF. This signal extends to nearly all 52 predictors examined, including population-free models unlikely to have been used in classification, probably reflecting shared evolutionary information. Classifications informed by computational evidence therefore cannot be treated as independent ground truth, and temporal holdouts cannot eliminate this circularity. To prevent a self-reinforcing feedback loop between prediction and classification, we recommend that ClinVar submissions routinely record the evidence criteria and computational tools supporting each classification in a standardised format, alongside greater use of classification-independent benchmarks.

## Background

The computational prediction of variant effects has a long history in human genetics. SIFT^1^ and PolyPhen^2^, two of the earliest methods, were built largely on the principle that amino acid substitutions at evolutionarily conserved positions are more likely to be damaging.

Machine learning subsequently enabled the integration of evolutionary conservation with structural, physicochemical and other features, while meta-predictors such as REVEL^3^ combined the outputs of existing tools. More recently, deep learning has substantially improved predictive performance, through unsupervised evolutionary models such as EVE^4^, protein language models such as ESM-1b^5^, and models building on protein structure prediction, such as AlphaMissense^6^. As methods have multiplied, so have the names used to describe them, including pathogenicity, deleteriousness, functional impact and in silico predictors. These methods are increasingly grouped under the broader term variant effect predictors (VEPs)^7^. Although most VEP development has focused on missense variants, the term also encompasses methods that score variants genome-wide, such as CADD^8^, and those that predict specific molecular consequences, such as altered splicing^9^.

A long-recognised challenge in the development and evaluation of VEPs is data circularity, where the same or related data are used for both training and assessment, inflating apparent performance^10^. Two main types have been described. Type I, or variant-level, circularity occurs when variants used to train or tune a predictor are later used to evaluate it. Type II, or gene-level, circularity arises when different variants from the same or homologous genes appear in both training and evaluation data. A predictor can then perform well by learning which genes tend to carry pathogenic variants, rather than by distinguishing between variants within them.

A third form, sometimes called type III circularity, has also been anticipated^10,11^. We refer to it as classification-level circularity, because it comes from how the labels themselves are assigned rather than from overlap between datasets. It occurs when VEP predictions contribute evidence to clinical variant classifications, and those classifications are later used to train, benchmark or calibrate VEPs. Predictors can therefore be evaluated on labels they helped generate, creating the potential for a self-reinforcing feedback loop between prediction and classification.

The direct impact of classification-level circularity was long assumed to be limited. The 2015 ACMG/AMP guidelines^12^, the standard framework for clinical variant classification, combine multiple lines of evidence, each assigned a strength from supporting to very strong (or stand-alone for benignity). Under these guidelines, computational predictions could provide only supporting evidence, through PP3 for pathogenicity and BP4 for benignity, and only when multiple tools agreed. The guidelines gave little direction on which tools to use or how to interpret their outputs, and supporting evidence alone has limited power to change a classification. Even so, it mattered in practice. In ClinGen expert panel curations made up to mid-2019, PP3 was applied to most pathogenic and likely pathogenic missense variants, and around one in six likely pathogenic classifications depended on it^13^. Predictions can also act indirectly, through ascertainment: variants with high scores may be prioritised for reporting, functional follow-up or curation. This enriches the pathogenic labels for variants that predictors already score highly, even when computational evidence was never formally applied.

As VEP performance improved^14^, capping computational evidence at the supporting level looked increasingly conservative. In 2022, the ClinGen Sequence Variant Interpretation (SVI) Working Group introduced a framework for calibrating individual VEP scores against ACMG/AMP evidence strengths and applied it to several widely used predictors^15^. Several reached strong evidence for pathogenicity, benignity or both, and the Working Group recommended that, for most situations, laboratories use a single genome-wide predictor reaching strong evidence for pathogenicity and moderate evidence for benignity.

The potential for VEPs to shape classifications directly is therefore now much greater. A single calibrated predictor can contribute up to strong evidence. Under the points-based evidence weighting expected in the forthcoming ACMG/AMP/CAP/ClinGen sequence variant classification standards (SVC v4.0)^16,17^, computational evidence combined with little else can move a variant out of the variant of uncertain significance (VUS) category. These developments make classification-level circularity an immediate concern. Here, we illustrate this emerging challenge by showing that VEP scores are already leaving a detectable signature in ClinVar^18^ classifications, and discuss how the community can address classification-level circularity before points-based standards become universal.

## A signature of computational evidence in ClinVar

We first considered four predictors, BayesDel_noAF^11^, MutPred2^19^, REVEL^3^ and VEST4^20^, which met the ClinGen SVI Working Group’s criterion for genome-wide single-tool use^15^ and are therefore most likely to be adopted as quantitative computational evidence. Each ClinVar submitter uses its own protocols, so the weight given to computational evidence, and which predictors are used, are likely to differ considerably between them. Any contribution of VEPs to classifications would be expected to appear unevenly, most strongly among submitters that have adopted specific predictors, and could be diluted across ClinVar as a whole. We therefore examined how mean scores for pathogenic (P), likely pathogenic (LP), likely benign (LB) and benign (B) missense variants changed over time for each of the largest ClinVar submitters, compared with all other submitters combined. To reduce confounding from changes in which variants are submitted, we restricted all analyses to rare variants (allele frequency <0.1%), since common variants tend to be identified and classified earlier, and considered LB and B variants only from genes with at least one P or LP missense variant in ClinVar. The latter restriction matters because some submitters classify large numbers of benign variants in genes with no established disease association, and changes in how many such variants they submit could otherwise drive apparent trends in mean scores.

For most submitters and classification categories, mean scores were broadly stable over time (Fig. 1a). One exception stood out: LB variants submitted by Ambry Genetics shifted abruptly towards more benign predictions for all four predictors between 2020 and 2021. Ambry LB variants now receive substantially more benign mean scores than B variants from Ambry or any other major submitter, for every predictor.

**Figure 1.**
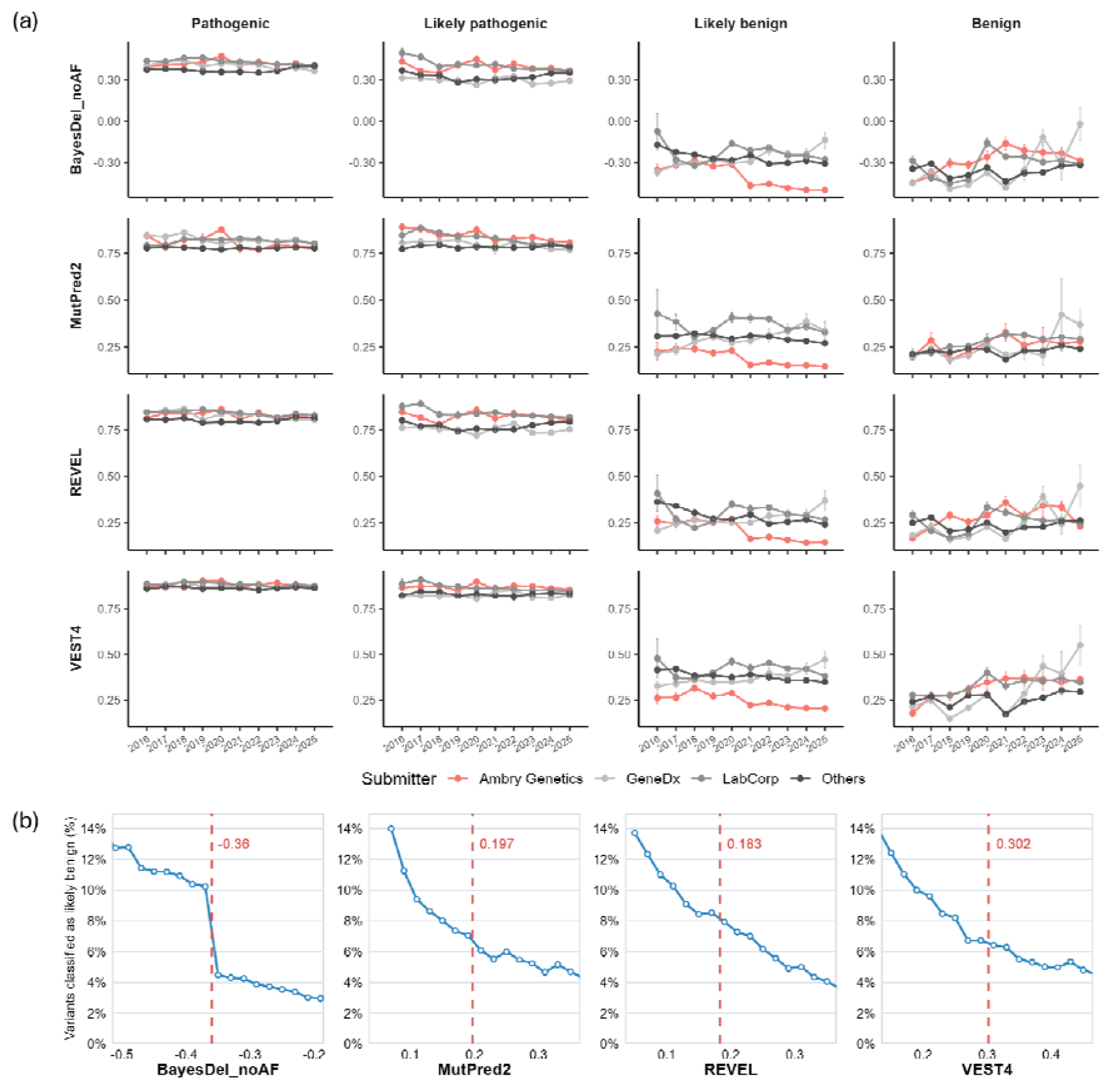
Signatures of computational evidence in ClinVar likely benign classifications. (a) Mean scores from four predictors (rows) for rare (<0.1%) disease-gene missense variants classified as pathogenic, likely pathogenic, likely benign or benign (columns), by year last evaluated (2016-2025). Lines show Ambry Genetics (red), GeneDx, LabCorp and all other submitters combined (“Others”). Higher scores indicate greater predicted pathogenicity. Error bars show SEM. (b) Percentage of Ambry Genetics rare missense variants classified as likely benign rather than VUS (LB/[LB + VUS]) as a function of predictor score, in bins of width 0.02, for variants last evaluated 2023-2025. Red dashed lines mark the BP4_Moderate thresholds^15^, with scores to the left providing moderate evidence for benignity.

Why would LB variants score as more benign than B variants, which represent considerably greater confidence in benignity? Some insight comes from Ambry’s 2023 Variant Classification Scheme^21^, a points-based framework in which LB requires a total of -2 or -3 points and B requires -4 points or lower. One possibility is the in silico criterion BP4, which contributes -1 to -3 points depending on prediction strength and could therefore reach LB on its own. ClinVar guidance states that classifications based solely on computational predictions are not appropriate for submission, but because submissions do not generally record the evidence behind each classification, those relying mainly on computational evidence cannot be identified. A second possibility is BP4_Ref, a fixed -3 point criterion applied when the variant amino acid matches the reference residue in other species, which is also sufficient on its own to reach LB. This follows the 2015 ACMG/AMP guidelines, which state that a variant amino acid present in multiple non-human mammalian species within an otherwise well-conserved region can be considered strong evidence for a benign interpretation^12^. Although BP4_Ref is not a computational prediction as such, it captures evolutionary information closely related to that used by many VEPs and would therefore be expected to correlate with their scores. However, while either alone can reach LB, neither alone can reach B. The LB category may therefore be disproportionately enriched for variants with strongly benign predictions, whereas B classifications must rely more heavily on additional evidence, such as population frequency or observations in unaffected individuals.

It is also possible that the shift in Fig. 1a partly reflects changes in the genes or variants Ambry assesses, which restricting to rare variants and disease genes controls for only partially. Direct use of VEPs, however, would leave a more specific signature. If Ambry assigns BP4 points as in the ClinGen points system, with moderate evidence worth -2, crossing the BP4_Moderate threshold would provide the -2 points needed for LB. If calibrated VEP scores were being used to assign BP4, we would therefore expect an abrupt increase in the proportion classified as LB rather than VUS at that threshold, beyond the gradual rise expected as scores become more benign. To test this, we examined Ambry rare missense variants around the BP4_Moderate threshold of each predictor, restricting to those last evaluated from 2023 onwards, the year Ambry published its scheme and the first full year after the ClinGen SVI calibrations (Fig. 1b). MutPred2, REVEL and VEST4 show only the gradual trend, whereas BayesDel_noAF shows a sharp step at its calibrated threshold of -0.36, with the proportion classified as LB increasing from approximately 4% to 10%. This strongly suggests that BayesDel_noAF was being used directly as computational evidence in Ambry’s recent classifications.

## Classification-level circularity extends beyond the predictors used in classification

Only a few missense VEPs have so far been calibrated for clinical use^15,22^. However, essentially all missense VEPs are correlated, because they draw on overlapping sources of information^23^. Variants classified partly on the basis of BayesDel_noAF scores will therefore tend to receive similar scores from correlated VEPs. The same applies to conservation-based criteria such as BP4_Ref, which do not involve any VEP directly but draw on the same underlying information.

To assess how widely this signal extends, we used 48 additional missense VEPs, covering a broad range of methodologies, to compare rare missense variants classified as LB by Ambry with those classified as LB by all other submitters (last evaluated 2023-2025). We quantified the differences using Cliff’s delta, a rank-based effect size that allows comparison across predictors with different score scales. Negative values indicate more benign scores for Ambry LB variants. Although other submitters’ LB classifications may also reflect computational or conservation-based evidence, this comparison indicates how strongly each VEP tracks the unusually benign scores characteristic of Ambry’s LB classifications.

BayesDel_noAF showed the largest negative effect size of any VEP (Fig. 2), consistent with its likely direct use in assigning BP4. However, all but one of the other VEPs also showed negative effect sizes. The sole exception, MAVERICK^24^, showed the greatest ancestry bias among 55 VEPs in our recent study^25^, while SNPred^26^ and MetaRNN^27^, which showed the smallest negative effects here, ranked second and third, respectively, for ancestry bias. This is consistent with these predictors relying heavily on ClinVar training labels and/or allele frequencies, which would favour more benign scores for variants classified using population evidence, as other submitters’ LB variants may more often be.

**Figure 2.**
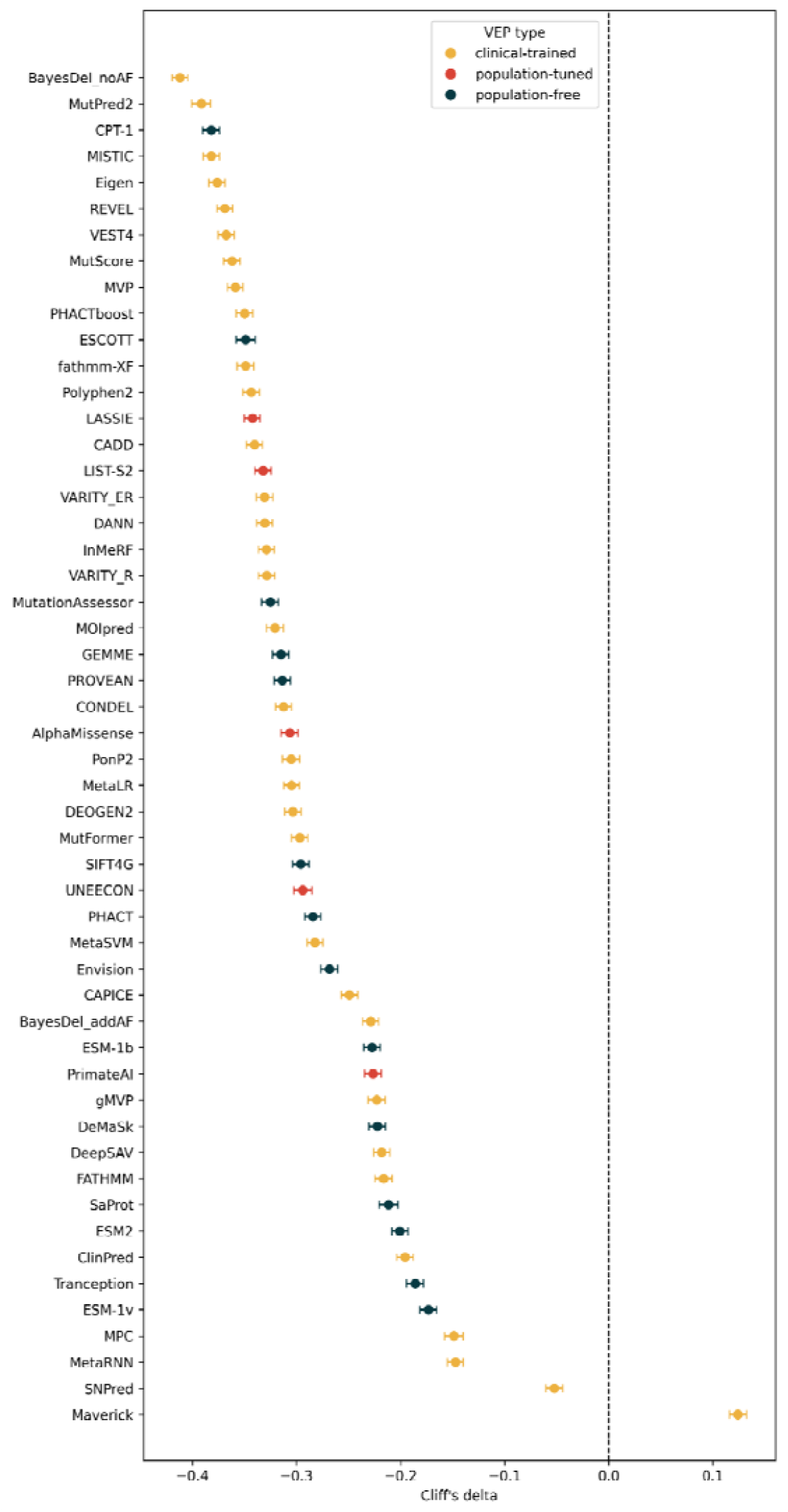
Predictor score differences between likely benign variants from Ambry Genetics and other submitters. Cliff’s delta comparing scores for rare (<0.1%) disease-gene LB variants from Ambry Genetics with those from all other submitters (last evaluated 2023-2025) for 52 different VEPs. Methods are coloured by type, as previously defined^14^: clinical-trained, trained directly or indirectly on clinically labelled human variants; population-tuned, tuned using human population variation but not clinical labels; and population-free, not trained or tuned using human population variation. Scores were oriented so that higher values indicate greater predicted pathogenicity. Negative values indicate lower (more benign) scores for Ambry. Error bars show 95% bootstrap confidence intervals (5,000 resamples).

By contrast, CPT-1^28^ showed the third-largest negative effect size, despite being a population-free model, not trained using clinical or population variant data, and being highly unlikely to have been used directly in variant classification. CPT-1 was also the best-performing missense VEP against MAVE data in our recent benchmarking study^14^, suggesting that its strong signal may reflect capturing much of the same evolutionary information underlying both BayesDel_noAF and conservation-based criteria such as BP4_Ref. More broadly, there was no clear relationship between effect size and the use of clinical or population variant data in VEP development, with substantial effects observed across all predictor categories.

## Consequences for training, benchmarking and calibration

ClinVar labels are among the most widely used resources for benchmarking predictors. If some of those labels were assigned partly on the basis of VEP predictions, performance estimates will tend to be inflated. The same problem applies to conservation-based criteria such as BP4_Ref. Observing the variant amino acid in other species is legitimate evidence of benignity, but the issue is whether it is independent of the predictors being assessed.

Since almost all VEPs draw on evolutionary conservation, evaluating them against labels assigned partly on this basis tests them on information they have already used, so agreement partly reflects shared inputs rather than independent predictive ability. Any errors in that information are also shared. For example, a pathogenic variant whose amino acid is tolerated in other species because of compensatory changes elsewhere in the protein^29^ may be classified as LB, scored as benign by VEPs, and counted as a correct prediction. The predictors used in classification will benefit most, but correlated predictors may also benefit indirectly.

This form of circularity cannot be avoided simply by using temporal holdouts, in which predictors are evaluated against variants classified after a particular date. Although this reduces overlap between training and evaluation data, newer classifications may rely more heavily on computational evidence. Lower performance on recently classified variants is often taken as a sign that older labels overstate predictor performance, but growing reliance on computational evidence could mask this drop or even reverse it.

Training on ClinVar raises closely related problems. Many supervised VEPs use ClinVar classifications as training labels, and where those classifications drew on computational evidence, models may partly learn to reproduce the predictions that shaped them, rather than independently distinguish pathogenic from benign variants. The risk is particularly direct for meta-predictors that include a VEP used in classification among their input features, since they may weight it heavily because it agrees with the training labels, regardless of its accuracy. Biases and errors in the original predictions can then be inherited by new models, whose outputs may themselves inform future classifications, creating a self-reinforcing cycle between prediction and classification.

Calibration presents a further concern. Because calibration estimates are derived from score distributions among classified variants, calibrations based on recent ClinVar data may partly reflect the computational evidence that contributed to those classifications, rather than independent evidence of pathogenicity. For genome-wide calibration, this problem can largely be avoided by using older variant sets that predate the widespread use of calibrated computational evidence and remain sufficiently large to calibrate new predictors^15^. However, VEP performance varies considerably across genes^30,31^ and there is growing interest in gene- and domain-specific calibration^32,33^. With classified variants spread across thousands of genes, historical datasets are often too sparse to support such analyses, creating pressure to use more recent classifications that are themselves more likely to incorporate computational evidence.

## Mitigating classification-level circularity

The extent of classification-level circularity in ClinVar-based datasets remains difficult to assess. Our aim is not to reconstruct or quantify all use of computational evidence in ClinVar, which may not be possible from public data, but to show that its influence exists and can be detected. While we found clear evidence of computational influence in one submitter and one classification category, this does not mean computational evidence has not contributed elsewhere. As the forthcoming SVC v4.0 will likely extend the points-based use of computational evidence across the field, these effects are likely to become more widespread, making it important to recognise and address them early.

We strongly support the increasing use of calibrated computational evidence in clinical variant classification. Many VEPs can justify stronger evidence than has traditionally been assigned to PP3 and BP4, with considerable potential to reduce the number of VUS, particularly among the growing number of rare variants lacking functional data. Assigning greater weight to computational evidence, as Ambry does, is consistent with ClinGen recommendations^15^, and Ambry’s points-based scheme is publicly available. A broader challenge for the field is that the precise computational evidence underlying classifications is not routinely documented: Ambry’s published scheme does not specify which predictors are used, and individual ClinVar submissions do not identify which predictions contributed to a classification. Our concern is not with the use of computational evidence in classification, but with a downstream consequence that is not yet widely appreciated: once VEP predictions have contributed to classifications, those classifications can no longer be treated as independent evidence for training, benchmarking or calibrating VEPs.

In the short term, the simplest precaution is to rely on older classifications that predate the widespread use of calibrated computational evidence, or, where more recent classifications are needed, to report results with and without them. Neither approach fully solves the problem. Older classifications may also reflect computational predictions, particularly at supporting strength, and effects on other categories or submitters may not produce a detectable signal in recent data. Restricting to older classifications also discards useful variants, a cost that falls most heavily on gene- and domain-specific analyses, where data are already sparse. Nevertheless, these approaches offer practical ways to limit, or at least assess, classification-level circularity until more systematic solutions are available.

A more durable solution is to record the evidence supporting each individual classification. The importance of documenting variant-specific evidence has long been recognised, particularly for resolving differences in classification between laboratories^34^. More recently, Dias et al.^35^ highlighted the circularity of evaluating models against labels derived from the same information, and recommended that ClinVar track the sources of evidence supporting each classification. Our results provide an empirical reason to act on that recommendation. ClinGen Variant Curation Expert Panels already document variant-specific criteria in the ClinGen Evidence Repository^36^, demonstrating that more detailed evidence recording is feasible. We suggest that ClinVar submissions routinely include the ACMG/AMP evidence codes and strengths applied to each variant, together with the predictor, version, score and threshold used for any computational evidence, ideally in a standardised, machine-readable format. This would allow users to identify classifications influenced by computational or conservation-based evidence, assess whether sufficient independent evidence remains to support them, and construct training, benchmarking and calibration datasets accordingly.

Such records would not capture indirect influences, such as variant prioritisation, but would make the direct contribution of VEPs transparent. Establishing this practice now would be far easier than reconstructing the evidence retrospectively once points-based classification becomes widespread.

Finally, alongside better evidence recording, greater use should be made of evidence independent of VEP-informed clinical classifications for developing, evaluating and calibrating predictors. Training VEPs without clinical labels avoids one important source of circularity, while functional measurements, de novo enrichment in developmental disorder cohorts and population-based comparisons provide complementary ways to assess their performance. Each has limitations, and independence is not automatic: functional evidence calibrated against the same clinical labels can inherit circularity, while population-based comparisons may overlap with information used to develop some predictors. Nevertheless, agreement across approaches with different sources of bias provides stronger support than agreement with ClinVar classifications alone. As VEPs become more influential in clinical classification, the evidence used to establish their reliability must remain independent of the classifications they help determine.

## Methods

### ClinVar data

Missense variant classifications were obtained from ClinVar, covering submissions with a date last evaluated up to the end of 2025. We used individual submitted records classified as pathogenic (P), likely pathogenic (LP), variant of uncertain significance (VUS), likely benign (LB) or benign (B). Ambry Genetics, GeneDx and LabCorp were analysed individually, and all remaining submitters were combined as “Others”. Each record was assigned to a year by its date last evaluated. Where a submitter had more than one record for the same variant, only the most recent was used. Variants classified by multiple submitters were retained for each submitter, so the same variant could appear in more than one submitter group.

### Variant filtering

Analyses were restricted to rare variants, defined as allele frequency <0.1% in gnomad v4.1, from genes with at least one P or LP missense variant in ClinVar, to limit the effect of changes in submissions from genes without an established disease association.

### VEP scores

VEP scores were obtained from the pipeline described in our previous study^14^, except for MutPred2, whose scores were downloaded from https://mutpred.mutdb.org/. All VEPs used in Pathak et al.^25^ with scores available for at least 70% of the disease-gene LB variants were included in the Fig. 2 comparison. Scores were oriented so that higher values indicate greater predicted pathogenicity. VEPs were categorised as clinical-trained, population-tuned or population-free as previously defined^14,25^.

### Threshold analysis

For Ambry rare missense variants classified as LB or VUS and last evaluated in 2023-2025, scores were grouped into bins of width 0.02. In each bin we calculated the percentage classified as LB, as LB/(LB + VUS). BP4_Moderate thresholds were taken from Pejaver et al.^15^: BayesDel_noAF -0.36, MutPred2 0.197, REVEL 0.183 and VEST4 0.302.

### Comparison across VEPs

For each VEP, we compared scores of rare disease-gene LB variants last evaluated in 2023-2025 from Ambry with those from all other submitters, using Cliff’s delta. 95% confidence intervals were estimated from 5,000 bootstrap resamples.

## Acknowledgements

This work was supported by funding from the Medical Research Council (MRC) Human Genetics Unit core grant (MC_UU_00035/9) and the Lister Institute Research Prize Fellowship.

